# The rebound response plays a role in the motion mechanisms and perception

**DOI:** 10.1101/2019.12.31.891580

**Authors:** Hadar Cohen-Duwek, Hedva Spitzer

## Abstract

Motion estimation is an essential ability for sighted animals to survive in their natural environment. Many anatomical and electrophysiological studies on low visual levels have been based on the classic pioneering HRC (Hassenstein & Reichaedt Correlator) computational model. The accumulated experimental findings, which have given rise to a debate in the current computational models regarding the interaction between the On and Off pathways. The previous algorithms were challenged to correctly predict physiological experiment results and the two types of motion: a) Phi motion, also termed apparent motion. b) Reverse-phi motion that is perceived when the image contrast flips during the rapid succession. We have developed a computational model supported by simulations, which for the first time leads to correct predictions of the behavioral motions (phi and reverse-phi), while considering separated On and Off pathways and is also in agreement with the relevant electrophysiological findings. This has been achieved through the well-known neuronal response: the rebound response or “Off response”. We suggest that the rebound response, which has not been taken into account in the previous models, is a key player in the motion mechanism, and its existence requires separation between the On and the Off pathways for correct motion interpretation. We furthermore suggest that the criterial reverse-phi effect is only an epiphenomenon of the rebound response for the visual system. The theoretical predictions are confirmed by a psychophysical experiment on human subjects. Our findings shed new light on the comprehensive role of the rebound response as a parsimonious spatiotemporal detector for motion and additional memory tasks, such as for stabilization and navigation.

## Introduction

The ability to detect motion is an essential quality for an animal’s survival (Clark et al. 2011; Arenz et al. 2017; Yang and Clandinin 2018), since it provides crucial information needed for exploring the external world. This information enables animals to perform tracking, navigation stabilization, and etc. A knowledge of the neuronal mechanisms and the structures of the motion detectors is critical to understand how an animal behaves in its complex world.

In general, animals, from mammals to insects, are sensitive to two types of motion. These were described by pioneering studies, as a common apparent or phi motion, and an illusory motion, the reverse-phi illusion (Anstis 1970). Phi motion requires at least two alternating stimuli, where the second stimulus, is slightly shifted spatially relative to the first stimulus. This configuration yields a perception of motion in the direction of the second stimulus. In contrast, for reverse-phi motion, the second stimulus is displayed with inverse contrast polarity (in addition to the shift in spatial location, as for phi motion). This yields a perception of motion in the opposite direction towards the first stimulus (Anstis 1970).

The low-level neuronal mechanisms and structures of motion detectors that can detect both these types of motions, have been the focus of numerous studies (Salazar-Gatzimas et al. 2018; Clark et al. 2011; Tuthill, Chiappe, and Reiser 2011; Theobald 2018; Luo-Li et al. 2018; Mo and Koch 2003; Leonhardt et al. 2017; Yang and Clandinin 2018). The pioneer motion study of Hassenstein and Reichardt (Von Hassenstein and Reichardt 1956), which used insects and has also been adapted to other animals, suggested a simple computational model (Alexander Borst 2000), the Hassenstein and Reichard Correlator (HRC), based on a delay and correlation mechanisms. However, the HRC was argued (Clark et al. 2011; Kuriki et al. 2008; Leonhardt et al. 2017) to be biologically implausible, due to the fact that the model requires a sign-correction of negative and positive signal multiplications in order to predict the reverse-phi phenomenon, and no neurons could fulfill such a function (Clark et al. 2011; Kuriki et al. 2008; Leonhardt et al. 2017).

In order to better understand the biological motion detection mechanisms, many electrophysiological, genetic, and behavioral studies have focused on the mammalian retina and the insect optical lobe (Alexander Borst and Helmstaedter 2015). The results of these studies revealed that changes in stimulus contrast polarity are processed through two motion pathways, one that detects an increase in stimulus brightness (an ‘‘ON’’ pathway) and a second that detects a decrease in stimulus brightness (an ‘‘OFF’’ pathway). These experimental results raise two important physiological questions, namely how the “ON” and “OFF” pathways interact with each other, and how such an interaction could explain the reverse-phi illusion.

With a consideration of the accumulated experimental knowledge, several computational models have been proposed in order to answer these questions (Clark et al. 2011; Mo and Koch 2003; Leonhardt et al. 2017, 2016; Salazar-Gatzimas et al. 2018). The models can be generally divided into two types, where the first assumes that directionality can be detected by discrete ON and OFF pathways (Leonhardt et al. 2017, 2016), while the second type assumes that there must be an interaction between the ON-OFF pathways, in order to correctly detect the direction of motion (Mo and Koch 2003; Salazar-Gatzimas et al. 2018; Clark et al. 2011).

The recent biological findings regarding the “OFF response” neuron, Mi9, which is connected to the ON pathway, provides supporting evidence for the models that assume the existence of interaction between the ON-OFF channels (Arenz et al. 2017). However, a more recent study demonstrated that this neuron (Mi9) does not influence the critical directional selectivity response of the directional selectivity cell, T4, in the fly’s optical lube, (Strother et al. 2017).

The second group of models, which assume separated ON and OFF pathways, can predict the reverse-phi phenomenon only by adding an additional DC component to the basic HRC structure of the model, (Leonhardt et al. 2017, 2016). Although some biological evidence for such a component has been found, it was only located before the processing of directional coding, and furthermore the directional selectivity (DS) cells (T4 and T5) apparently do not carry a DC signal (Leonhardt et al. 2017). This finding is supported by the observation that T4 and T5 DS cells have precisely polarity-specific edge responses (Maisak et al. 2013).

Consequently, there is no unequivocal biological evidence to fully support the assumptions of models involving an interaction between the ON and the OFF pathways or for models that describe separated pathways but require a DC component. We therefore examined the possibility that there is a different biological component that can be integrated into the basic HRC model to predict both the phi and the reverse-phi phenomena, and accord with the various physiological findings.

In this paper, we present a computational model that predicts both phi and reverse-phi, by using the well-known “rebound” biological response, which signals the “turning – off” of a stimulus (Enroth-Cugell and Jones 1963; D. Hubel 1995; Kuffler, Nicholls, and Martin 1984).

## Results

### model

Our suggested model provides a novel approach to the elucidation of motion detection mechanisms, even though the basic structure of the model’s architecture is similar to previous models (besides accounting for the cardinal neuronal response). Our model, as well as the previous motion models, are small variants of the classical HRC model of Hassenstein and Reichardt (Von Hassenstein and Reichardt 1956; Mo and Koch 2003; Leonhardt et al. 2017, 2016; Salazar-Gatzimas et al. 2018; Adelson et al. 1985) and share a common architecture (Clark et al. 2011; Mo and Koch 2003; Leonhardt et al. 2017, 2016; Salazar-Gatzimas et al. 2018; Tuthill, Chiappe, and Reiser 2011; Yang and Clandinin 2018), consisting of the following layers: 1) the photoreceptor layer, Fig 1 (a). 2) a rectification layer (Reiff et al. 2010), which converts the signal to ON and OFF pathways, (e.g., L1/L2 in fly’s lamina layers and retinal bipolar cells in mammalians), Fig 1(b). 3) a delay layer, Fig 1 (c). 4) the directional selectivity cells layer, which detects motion in a specific direction (T4/T5 in flies, and SAC cells in mammalians), Fig 1 (d).

**Fig 1:**
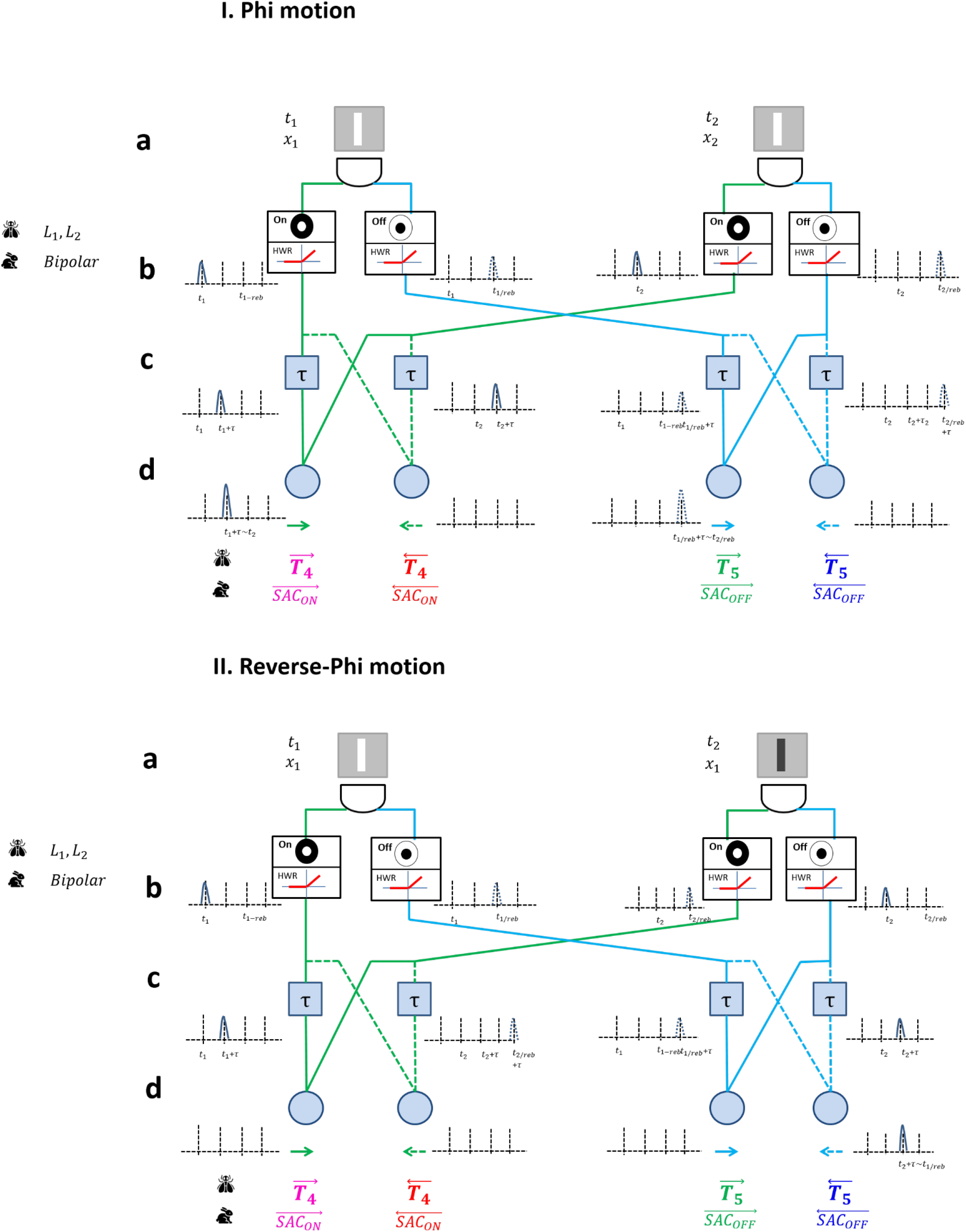
The Model predicts both phi and reverse-phi motion (phi (I) and reverse phi (II)) for light and dark bars. The model demonstrates in a simple way its predictions through the illustration of two photo-receptor units and their appropriate wiring pathways. The left unit represents the location of the first stimulus (x1) and time (t1), while the right unit represents the second location(x2) and time (t2). A directional selectivity (DS) response can be achieved through the idea that a response from the first unit is delayed through a delay component (tau), and this response can be superimposed with a response from the second unit at a shifted location and at a later time. The DS response has to be followed by the rules of connectivity, which have been found experimentally, i.e., On (green lines) and Off (blue lines) pathway responses are processed separately. Phi motion: A stimulus provided by a light bar is moving from the left location to the right location (a). Two types of center-surround cells, On-center and Off-center, receive input from the same photoreceptor: The light bar simultaneously activates the On-center cell and inhibits the Off-cell (b). These signals are also half-wave-rectified (HWR), and thus, only transmit positive values for the responses (b). The output from On-cell, at the first location, is then temporally delayed by the delay component, τ (c). When the light bar stimulus is presented to the second photo-receptor, it also activates the On-center cell and inhibits the Off-center cell. Both the delayed signal from the first location and the signal from the second location arrive at the directional selectivity cell (T4) simultaneously and activate it, while the other DS cells remain inactivated. Reverse-phi motion: in this type of motion the stimulus is presented at the second location with an opposite polarity, therefore, the On- pathways are not activated from both photoreceptor units. The DS cells are only activated when the rebound response from the Off cell (x1) (after the stimulus is turned off at this location) in the first location is superimposed on the delayed Off cell response from the second location(x2). In this case, when a light bar and then a dark bar are presented, a leftward T5 DS cell is activated.

Our model also consists of the above architecture, Fig 1, but in contrast to previous studies that added components to the HRC basic model to enable the prediction of the reverse-phi motion (Salazar-Gatzimas et al. 2016; Clark et al. 2011; Leonhardt et al. 2017), we have implemented a well-known neuronal response i.e., the rebound response, and combined it in the motion detector model. This type of response yields a neuronal excitatory response after an inhibitory stimulus is turned-off (stimulus offset) (Enroth-Cugell and Jones 1963; D. Hubel 1995; Kuffler, Nicholls, and Martin 1984). Significantly, the rebound response has been neglected by all the previous motion models.

According to our model (and the other models), a phi movement of a light (or dark) bar, Fig 1 (I), is perceived as moving from left to right (rightward motion), as detailed in Fig 1 (I). This motion directionality is detected when both the delayed signal from the left location (*x*_1_) and the signal from the right location (*x*_2_) arrive at the motion detector layer nearly simultaneously, Fig 1 (d), and excite a motion detector cell, Fig 1 (d) to identify a rightward motion (solid green lines, Fig 1). Our model, thus predicts the correct phi-motion direction in a similar way to many other variants of the HRC model (Clark et al. 2011; Mo and Koch 2003; Leonhardt et al. 2017, 2016; Salazar-Gatzimas et al. 2018; Tuthill, Chiappe, and Reiser 2011; Yang and Clandinin 2018).

The directionality prediction of the rightward reverse-phi movement of a light bar is derived from the simultaneous arrival of the rebound response, at *x*_1_, with the response of the Off cell, at *x*_2_, to the dark bar. The rebound response at *x*_1_ is caused by turning off the light bar at the Off cell. These superimposed responses arrive almost simultaneously at the motion detector, *T*_5_, which then indicates motion directed towards the left. Consequently, an opposite motion detector, i.e., of the leftward motion, is activated in the reverse-phi illusion. Our model, thus, predicts the perceived motion, e.g., a rightward reverse-phi movement of a light (or dark) bar is perceived as moving in the opposite direction (from right to left), Fig 1 (II).

Since the rebound response is very common in the visual system and always occurs after an inhibitory response (Kuffler, Nicholls, and Martin 1984) temporal edge, many mechanisms and particularly those connected to motion have had to take this type of response into account in their mechanisms and therefore in their models. However, considering the described occurrence of a rebound response, together with a scenario in which the On and Off pathways interact (are wired together) can lead to exciting a “wrong” motion detector. The “wrong” motion detection can be demonstrated, for example, as follows: the signal of the rebound response to a light bar stimulus from the left location (x1), which is superimposed simultaneously on the On response that arrives from the second location (x2), will activate an opposite DS cell (opposite to the direction of the real movement).

### Simulations and experiment

The model and its predictions have been tested by both simulations, and a psychophysical experiment, which examined whether there is a simple way to abolish the reverse-phi effect and thus to shed more light on this enigmatic effect.

### Simulations

Fig 1 presents the model simulations as schematic block diagrams for phi and reverse-phi, types of motion. We applied an atrophied version of an optical flow algorithm (i.e. correlation calculation (Beauchemin and Barron 1995)), in order to obtain the model’s predictions for the apparent motion of a bright or dark bars, as the visual stimuli (Methods). The two types of stimuli (dark and light bars) are presented on a gray background (Fig 2), and are detected through a matrix of On and Off center-surround receptive fields. These receptive fields map the neuronal net and contain a complete overlap of the On- and the Off receptive fields, at the locations of the left and right “motion” directions, Fig 1. For the sake of simplifying the computations of the receptive field matrix and the center-surround receptive fields in the fly’s optic lobe, we only subtracted the background (which is used as the surround region) from each pixel (which is used as the center) (Freifeld et al. 2013).

**Fig 2:**
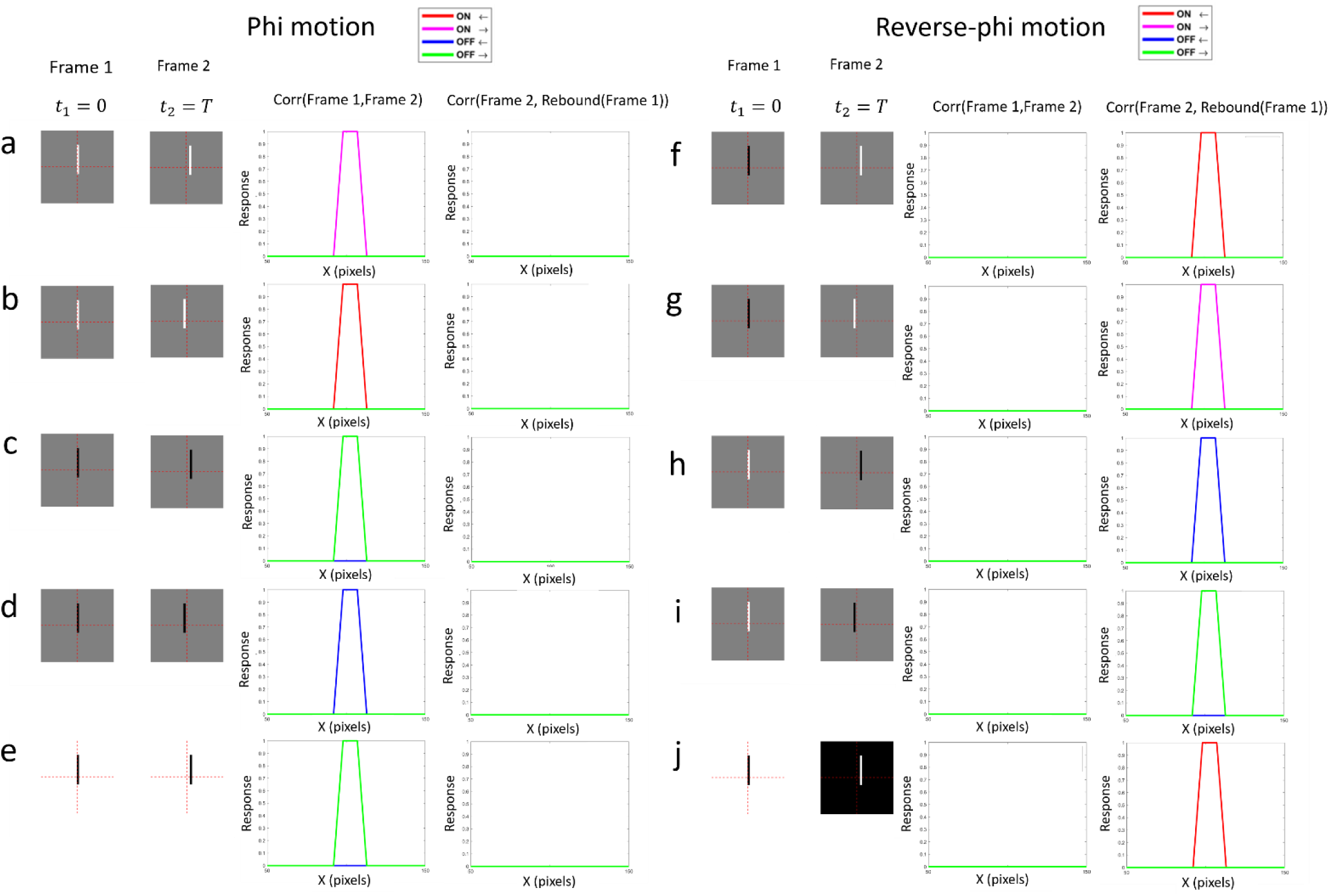
The simulated cross-correlation responses obtained from the phi and reverse-phi motion stimuli. The stimuli are presented in the framework of apparent motion, i.e., with the images presented in rapid succession. The first cross-correlation (left response column) was calculated between Frame 1 (the stimuli at time t1) and Frame 2 (the stimuli at time t2), for each type of stimulus, separately. The second cross-correlation (right response column) was calculated between Frame 2 and the rebound response of the stimulus at t1 (rebound responses of Frame 1). The figure presents 10 combinations of light and dark bars moving to the right or left, in both phi and reverse phi types of motion. Phi motion: This type motion is illustrated through a light bar stimulus “moving” to the right (a) or to the left (b); while a dark bar stimulus was presented as “moving” to the right (c) or to the left (d). A dark bar is presented as “moving” from left to right (e), on a white background. Reverse-phi: A dark bar is presented as moving from left to right (g) and right to left (g). A light bar is presented as “moving” from left to right (h) and from right to left (i). A dark bar is presented as “moving” from left to right (j) on a white background.

Note that in accordance with reports on the pathways in the visual system of flies (Alexander Borst and Helmstaedter 2015) and previous computational studies (Salazar-Gatzimas et al. 2018; Clark et al. 2011; Tuthill, Chiappe, and Reiser 2011; Theobald 2018; Luo-Li et al. 2018; Mo and Koch 2003; Leonhardt et al. 2017; Yang and Clandinin 2018), we considered only the simple case of two directions of motion although the visual system of flies can sense four directions of motion in the Labula visual layer (Alexander Borst and Helmstaedter 2015).

The model simulations indicated that a phi motion with a light stimulus, Fig 2(a-b) caused the On DS cells to provide a significant response, while a phi motion with a dark stimulus, Fig 2 (c-d) caused the Off DS cells to respond significantly. In the reverse-phi motion, however, the responses were different. With a dark bar in the *x*_1_ location (frame 1) and a light bar in the *x*_2_ location (frame 2, Fig 2 (f-g)), the activated DS cells belonged to the On type. In the opposite case, when the bar in the *x*_1_ location was a dark bar (frame 1) and the bar in *x*_2_ location was a light bar (frame 2), the activated DS cells were Off cells. Consequently, the model simulations, Fig 2, succeed in demonstrating that the suggested model with its simple architecture can lead to the correct predictions of both phi and reverse-phi, across the different stimuli polarities, and the two different directions of motion, Fig 2.

The correct predictions of the directions of the reverse-phi stimuli are obtained due to the responses to “turning off” the stimuli (i.e., rebound) from the receptive field of the first location (Fig 2, Frame 1), which coincide with the response to the stimulus with opposite polarity from the response of the second location, (Fig 2, Frame 2). The results clearly show that significant responses are obtained at the turning-on responses for the phi motion, Fig 2 (a-e), while significant responses are obtained at the turning-off responses for the reverse-phi movement, Fig 2 (f-j). Note that the reverse-phi motion therefore occurs at a slower rate or is temporally delayed.

### Psychophysical experiment

Our motion model is based on separated On and Off pathways with the rebound effect. Even though the rebound effect always exists, it is considered here only with respect to its role in reverse-phi motion perception.

It has been suggested that the mechanism for the aftereffect is the rebound response (Grossberg 1976; Francis 2010; Zaidi et al. 2012; Cohen-Duwek and Spitzer 2018). Earlier studies showed that a noise mask can abolish the aftereffect phenomenon (Haber and Standing 1970; Felsten and Wasserman 1980). We assumed that if both aftereffect and reverse-phi effect are derived from the same rebound mechanism, then the noise mask should abolish the revere-phi effect, as well as the aftereffect. This mask noise technique was used in the psychophysical experiment for both the phi and reverse-phi stimuli, through introducing a noise mask after turning off the first stimulus, and before activating the second stimulus.

The psychophysical results, Fig 3, demonstrate that all four observers obtained high proportions of correct responses scores with both phi and phi with noise stimuli, with close to the maximum proportion of correct responses (1). However, the observers were less successful with the reverse-phi stimuli. Interestingly, the proportion of correct responses by all the four observers significantly increased under the masked reverse-phi stimulus condition, in comparison to the correct responses to reverse-phi stimuli without the mask, Fig 3.

**Fig 3:**
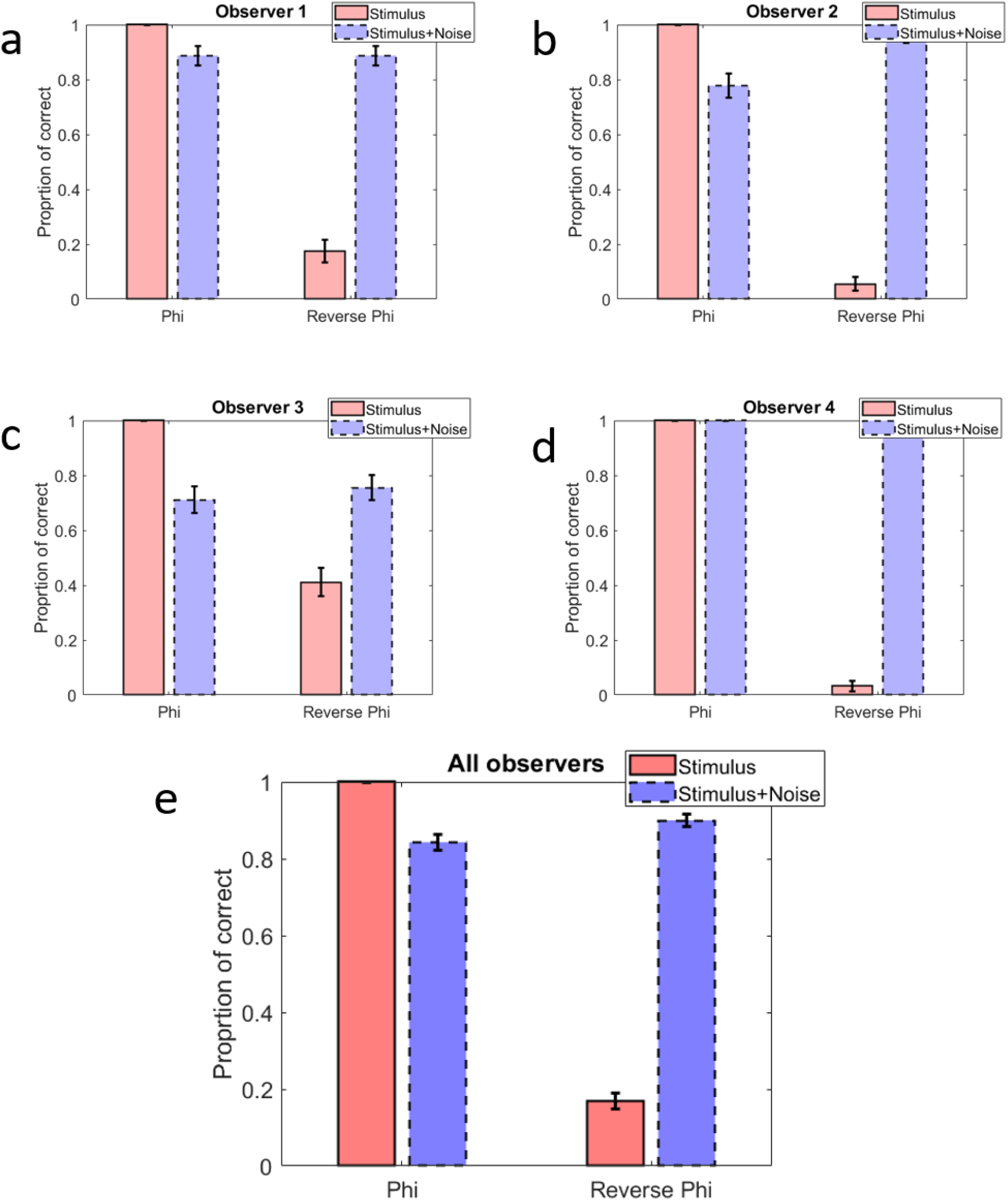
presents the proportion of correct responses of 4 observers to the directional motion of phi, and reverse phi and the responses of these two types of motions with a noise mask. This type of basic stimulus has been applied for all types of stimulus (phi, masked phi, reverse phi, masked reverse phi). The observers answered the question as to whether the set of circles moved, clockwise or counterclockwise, as part of two forced-choice tasks. The most prominent result was that the noise mask “succeeded” in reversing the low performance of detecting the correct direction of apparent motion in response to the reverse-phi stimulus. This pattern of response was consistent and was statistically significant across all the four observers, even though they displayed variability in their level of correct responses, mainly to the stimuli with the noise mask.

The psychophysical results indicate that the addition of a noise mask frames significantly improved the proportion of correct responses in the reverse-phi stimulus type and brought the scores close to the level obtained with the masked phi stimuli. It should be noted that the addition of noise mask frames, had only a minor influence on the responses to the phi type of stimulus type. The results for phi and reverse-phi stimuli, suggest that the introduction of a noise mask cancels the reverse-phi illusion, as has been assumed if this model is based on the role of the rebound response.

## Discussion

We suggest here a new motion model that can predict both phi and reverse-phi motion effects, assuming separated On and Off pathways and using only known and proven neuronal responses and mechanisms. The model applies a critical component, which is a fundamental neuronal mechanism, i.e., the rebound response (Schwartz, Kandel, and Jessell 1991; D. Hubel 1995). This response is an excitatory response that occurs when an inhibitory stimulus is turned off (Schwartz, Kandel, and Jessell 1991) (Schwartz, Kandel, and Jessell 1991; D. Hubel 1995). Applying this mechanism to a version of the HRC motion model, with separated On and Off pathways, as found physiologically (Bausenwein, Dittrich, and Fischbach 1992; Maisak et al. 2013; Kim et al. 2014), enables us to predict both phi and reverse-phi motions correctly, without the need for any additional components (such as additional DC components and different weights for the different On and Off pathways), as have been suggested previously (Salazar-Gatzimas et al. 2018; Clark et al. 2011; Mo and Koch 2003; Leonhardt et al. 2017, 2016).

In order to assess whether the rebound response is crucial to the perception of motion and assess the importance for the prediction for both the phi and reverse-phi movement, we conducted a psychophysical experiment that disrupts the mechanism of the rebound response and therefore abolishes the aftereffect (Haber and Standing 1970). For this purpose, we used a noise mask (Haber and Standing 1970; Felsten and Wasserman 1980), and tested whether abrogating the aftereffect affected the reverse-phi perception. The results presented in Fig 3, demonstrated that the reverse-phi illusion (aftereffect mechanism) was indeed abolished by the noise mask, and therefore, it indicates that the aftereffect, which is resulted by the rebound mechanism (Grossberg 1976; Francis 2010; Zaidi et al. 2012; Cohen-Duwek and Spitzer 2018), is essential for perception of the reverse-phi motion.

The existence of the rebound response in the retina/insect’s optic lobe, has been proven by many electrophysiological studies on insects and mammals, after its presence was previously well established in cats and primates (Kuffler, Nicholls, and Martin 1984). The rebound response has been found, for example, in the Lamina Monopolar cells (LMS) of insects (Alexander Borst and Helmstaedter 2015; Laughlin, Howard, and Blakeslee 1987), and in the retinal bipolar and ganglion cells of mammals (Mangel 1991). The rebound response has also been found, on the other hand, in the higher visual areas of mammals such as the v1, v2 MT cells of monkeys (D. H. Hubel and Wiesel 1962; Livingstone, Pack, and Born 2001; Kuffler, Nicholls, and Martin 1984). In addition, the rebound response has been found in the auditory and the somatosensory systems (Rajaram et al. 2019). These findings, thus, imply that the rebound response is a fundamental component of an animal’s neuronal system. This prevalence of the rebound response strongly supports the need to integrate it as a component in a computational model.

### Previous computational models

We would like to scrutinize here the previous models from the two categories related to the interactions between the On and Off pathways: (a) with overlapping ON-OFF channels (Salazar-Gatzimas et al. 2018; Clark et al. 2011; Mo and Koch 2003) and (b) with separated ON-OFF channels, in the light of recent experimental studies (Leonhardt et al. 2017, 2016).

### Overlapping channels

A number of computational studies (Salazar-Gatzimas et al. 2018; Clark et al. 2011; Mo and Koch 2003) have suggested models with overlapping channels, but with the implementation through weighted interactions between the ON and OFF pathways. According to these models, each input pathway connected to the directional selectivity cells (T4 - On pathway or T5 - Off pathway), has a different weight. These weighted channels enable the computational models to predict both phi and reverse-phi effects, and to show asymmetric properties of the ON and the Off pathways, in accordance with experimental results (Clark et al. 2011; Leonhardt et al. 2016).

A recent study (Salazar-Gatzimas et al. 2018) provided evidence that supports the overlapping On and Off channel models, through the findings that the T4 and T5 cells respond to both phi and reverse-phi stimuli. It should be emphasized that the reverse-phi stimuli always contain both polarities of dark and bright stimuli. The additional evidence, in favor of the On and Off interaction, came from the findings of the “Off-response” neuron, Mi9, as an input to the T4 cell (Arenz et al. 2017). The authors (Salazar-Gatzimas et al. 2018), consequently, concluded that the computational mechanism has to combine the On and the Off pathways.

We argue that this conclusion is not necessarily correct for two reasons: the first being that that there is an additional explanation with separated ON-OFF channels that can explain their results. We have shown in the Model Simulations, Fig 2, that indeed both the phi and the reverse-phi can be predicted by our computational model, Fig 1, which assumes separated On and Off channels. This prediction can be demonstrated in the model’s simulation, Fig 2, where each different type of On or Off DS cells respond to both phi and reverse-phi stimuli, even though the On and the Off channels are separately wired.

The second reason to doubt the ON-OFF interaction conclusion is derived from new findings (Arenz et al. 2017) showing that silencing the “Off response” input neuron, Mi9, (belonging to the On pathway) does not influence the directional selectivity response of the DS cell, T4 (Strother et al. 2017).

### Separated channels

Computational models assuming separated On Off pathways (Leonhardt et al. 2017, 2016) are based on experimental findings (Bausenwein, Dittrich, and Fischbach 1992; Maisak et al. 2013; Kim et al. 2014), such as studies that showed that genetically blocking the lamina monopolar cells L1 (the beginning of the ON pathway) suppresses the ON-ON responses, but leaves the OFF-OFF responses intact (Maisak et al. 2013). As a corollary, blocking the output from the lamina monopolar cells L2 (the earlier stage of the OFF pathway) was shown to leave the ON-ON responses intact (Maisak et al. 2013). These results imply that there are two separated physiological ON and OFF pathways (Alexander Borst and Helmstaedter 2015; Mauss et al. 2017). Our model is in agreement with this implication (Maisak et al. 2013).

However, since a simple HRC model with separated ON and OFF pathways cannot predict the reverse-phi illusion (Tuthill, Chiappe, and Reiser 2011) it was necessary to add a DC component (Leonhardt et al. 2017, 2016). With this addition, the model does succeed in predicting the reverse-phi effect when applied through modeling a neuronal response that contains a combination of transient (High Pass) and tonic (DC) temporal components (Leonhardt et al. 2017).

Electrophysiological studies, in the fly’s optic lube, supported the use of the DC component, by demonstrating the existence/presence of the tonic response in the fly medulla (Serbe et al. 2016; Arenz et al. 2017; A. Borst and Bahde 1986; Behnia et al. 2014). However, the implications of these results might be disputed, since the tonic effect has not been found in the critical DS cells (T4 and T5), (Leonhardt et al. 2017; Fisher, Silies, and Clandinin 2015; Maisak et al. 2013; Salazar-Gatzimas et al. 2016).

Our model is in agreement with the idea of separated ON and OFF pathways, and with the physiological findings of polarity-specific edge responses of the DS cells, T4, and T5 neurons (Leonhardt et al. 2017), while not requiring any type of DC component. It should be noted that in our simplified model simulations, we neglect the temporal shape of the neural responses, as we only consider the temporal properties by the “turning on” and the “turning off” (rebound) responses. The crucial mechanism that explains the reverse-phi in our model is the rebound response and not the temporal shape of the neuronal response.

While experimental biological and electrophysiological results might be considered to challenge previous models (Salazar-Gatzimas et al. 2018; Clark et al. 2011; Mo and Koch 2003; Leonhardt et al. 2017, 2016), our model has not yet been tested against physiological observations. We propose at this stage a novel psychophysical experiment on human observers that has the potential to confirm or refute our suggested model.

We were able to abolish the reverse-phi perception, for all four observers (Fig 3) by applying a simple noise mask that eliminated the rebound response in the experiment (Fig 3), we found that the mask abolishes the reverse-phi perception. This psychophysical result therefore provides strong support for our underlying assumption that the rebound effect is the root cause of both the aftereffect and the reverse-phi effect, i.e., canceling the rebound response cancels the reverse-phi illusion. We believe that all the other models should also be confronted with our new psychophysical effect, and challenged to predict the existence or abolition of the reverse phi effect by the addition of a noise mask.

Since the previous models (Salazar-Gatzimas et al. 2018; Clark et al. 2011; Mo and Koch 2003; Leonhardt et al. 2017, 2016) do not contain components that are affected by such factors as a noise mask, we assume that they will not be able to predict the cancellation of the reverse-phi effect by the mask, but appreciate that it has to be tested directly.

Indeed, the reverse-phi illusion has found much use as one of the validation criteria for motion models. This raises the question of whether there is a general functional role of the reverse-phi with computational/other benefits. We ask here whether this type of effect has a functional role in a natural environment and in the motion mechanism itself, or is rather just a side-effect of the motion mechanism. A recent study (Salazar-Gatzimas, Agrochao, Fitzgerald, Correspondence, & Clark, (2018)) suggested that the reverse-phi effect is an essential part of the motion mechanism. The reasoning was that such an effect occurs in natural environments when the edges of an object switch polarity as an animal moves. The authors of the study (Salazar-Gatzimas, Agrochao, Fitzgerald, Correspondence, & Clark, (2018)) suggested, based on their computational model, that the reverse-phi effect plays a role in the animal’s ability to precisely predict the velocity of the object.

We, however, we do not agree with this concept (Salazar-Gatzimas, Agrochao, Fitzgerald, Correspondence, & Clark, (2018)), since it is not common for an object to abruptly switch polarity in a natural environment. We do appreciate that the edges of an object might sometimes appear to switch polarity, for example, if there are repeated instances of an object in the scene (e.g., tree boulevard). However, such scenarios are rare and we do not see any biological advantage in obtaining a perceived image of an object apparently moving in the opposite direction (a response to an illusory direction).

### The role of the rebound response

In contrast, to the previous considerations, the rebound mechanism can predict additional motion illusions, such as “motion aftereffect” or “waterfall illusion” (Barlow and Hill 1963), which are derived from real motion. The perception of an object moving in the opposite direction (Barlow and Hill 1963) as part of the motion aftereffect phenomenon after turning off the stimulus, might be explained as a rebound response of the Lobula Plate Tangential Cell (LPTC), which receives inputs from T4 and T5 cells. Support for this hypothesis provided by physiological results showing that LPTCs are depolarized most strongly in response to a specific directional stimulus and hyperpolarize when the direction of motion is reversed (A Borst and Haag 1991). The motion direction pathway signaling direction through hyperpolarization (to the non-preferred stimulus) might be switched when the stimulus is turned off and an excitatory response (the rebound response) is produced (after inhibition).

Simulations with the model support the idea that both types of motion can be accurately predicted by considering separated On and Off pathways. Thus, our model strongly supports the separationist side of the debate as to whether the early motion mechanism originated from mixed or separate On and Off mechanisms. Parallel On and Off pathways can be advantageous for the visual system, beside the role in motion mechanisms. For example, processing information through two independent pathways can provide a certain redundancy which can be used to validate system decisions (Cover and Thomas 2012).

The model succeeds in predicting both phi and the reverse-phi motion with discrete On and Off pathways through the integration of the rebound response. Physiologically, the rebound response is an essential component involved of low level motion mechanisms, as found in the retina or flies’ optical lobe as well as in cortical levels. Support for the significance of the rebound component can be seen in the wide prevalence of the phenomenon in different visual system stages and areas. Recently, studies have also implicated the rebound response in the issue of saccade-based termination (Niemeyer and Paradiso 2018) and object motion versus motion from eye movements (Troncoso et al. 2015). The function of the rebound response in eye movements might also play a role in other mechanisms, such as visual stabilization and navigation. In addition, the rebound response can be used by the visual system as a tool for temporal memory storage and, thus, to overcome the tendency of certain neurons to signal a transient response and, therefore, to transmit only or mainly, changes in time. Given the wide variety of roles and locations in which this effect is present, it would appear to be very important to further investigate the mechanism of the rebound response and its connection to other sensory systems. In addition, modeling the separated On and Off pathways together with the integration of the rebound response, serving as a spatio-temporal detector, should provide information beyond their connection to motion encoding. This could be used, for example, to create two independent channels that can be used to analyze decisions made under ambiguous conditions such as often occur in the 3D world, with variable illumination, occlusion problems and etc.

## Methods

### Model

The model proposed here is based on a variation of the HRC model (Franceschini, Riehle, and Le Nestour 1989). The architecture of our model requires separately wired ON and OFF pathways, Fig 1. The main building blocks of the model are: (A) The input: the apparent moving stimuli (light or dark bars), Fig 1(a), which are fed into the photoreceptors. (B) On and Off opponent receptive fields, Fig 1(b). (C) The temporal delay Fig 1(c). (D) Direction selectivity (DS) cells, Fig 1(d).

The model is a correlative model, which performs a correlation between two signals that are separated both in time and space domains. This type of correlation is similar to the common HRC model and its variants (Von Hassenstein and Reichardt 1956), Fig 1(d).

### Input: The moving stimuli

The stimuli consist of light or dark bars that move from left to right or vice versa. In the phi motion stimuli, a light or dark bar moves, while in the reverse-phi stimuli a light or dark bar moves and then changes its polarity at the second spatial location (i.e., a dark bar becomes a light bar at a later time), Fig 1 I(a) and Fig 1 II(a). The direction of the moving bar is determined according to the order of the temporal appearance of the bar at the two locations. If the bar is displayed first at the right side, and then at the left side, the real direction is leftward, and vice versa to the other direction: rightward. These stimuli are translated to neuronal responses at the level of the photoreceptors. The size of each stimulus is 200×200 pixels and the size of the bar in each stimulus is 81 pixels in height and 6 pixels in width. The bar is shifted by 9 pixels between Frame 1 and Frame 2, Fig 2.

### The On and the Off Receptive fields

The second computational stage involves the On and Off receptive fields, Fig 1 I(b) and Fig 1 II(b). The model refers to On and the Off receptive fields of L1 and L2 cells in flies or in the Bipolar cells of mammals, both of which possess a center-surround opponent receptive field structure, (Freifeld et al. 2013; Strother et al. 2017; Arenz et al. 2017). The response of center-surround opponent receptive fields is half-wave rectified in the model, due to electrophysiological findings in both fly L1, L2, and mammalian bipolar cells (Reiff et al. 2010). These cells are the first chain in the separated On and Off pathways. We took into account here the classical center-surround receptive field with an annular surround shape. This receptive field is mathematically expressed as Laplacian of Gaussian or Difference of Gaussian. However, for the sake of simplicity, we implemented the center-surround opponency by subtracting the intensity of the image background, from each pixel’s intensity to give the surround region. We calculated the On or the Off receptive field response (*I*_*t,On*_) by subtracting the background (of the light or dark bar) from their center response, and then rectified it. This has been done across each pixel of the image, Fig 2, (Reiff et al. 2010). The half-wave-rectified operation was done through a max function, Equation (1). In order to calculate the Off channel (*I*_*t,Off*_), we performed the opposite subtraction, i.e., subtracting the pixel’s intensity from the intensity of the background, and half-wave-rectified the result, Equation (2).

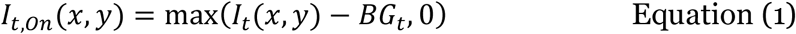

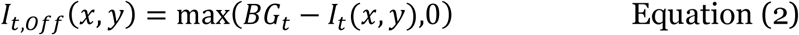

where *I*_*t*_ is the intensity of the image (which is composed of a bar stimulus and a background, Frames 1-2 in Fig 2) at time, t = {0,T} (time t = 0 or t = T, respectively, Fig 2). *BG* is the background of the image (BG = 0.5 in the stimuli (a-d) and (f-i), BG = 1 in the stimulus (e) and in the first frame of the stimulus (j), and BG = 0 in the second frame of the stimulus (j), Fig 2).

### The Rebound Response

The rebound response is first applied in our motion model, Equation (3-4). It is an excitatory neuronal response that occurs after turning off an inhibitory stimulus (Kuffler, Nicholls, and Martin 1984). This classical response was found in many stages in the visual system including the Bipolar cells in the mammalian retina, and L1 and L2 cells in the fly optical lobe (Laughlin, Howard, and Blakeslee 1987). We suggest here that this type of response is essential for understanding a mechanism that is related to the basic motion effects in the optic lobe or the retina.

Even though the model for the rebound response, including its properties and parameters, have been suggested previously (Enroth-Cugell and Jones 1963; Spitzer, Almon, and Sandler 1993; Sherman, Almon, and Spitzer 1994), we here refer only to its occurrence in space and time domains. For the sake of simplicity, we calculate the rebound response as the inverse response of the half-wave rectified On or the Off receptive field responses, as follows:

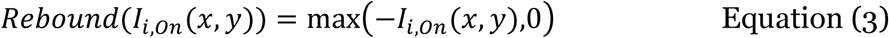

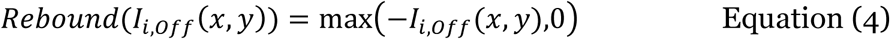

### Temporal Separation (Delay)

The delay component is an essential component in our model, as well as in the HRC model (Von Hassenstein and Reichardt 1956) and all its variants (Salazar-Gatzimas et al. 2018; Clark et al. 2011; Mo and Koch 2003; Leonhardt et al. 2017, 2016). It is reasonable to assume that the visual system has many motion detectors that vary in their spatial distance and temporal properties, including the magnitude of the delay and the latency of the rebound responses. In our model, for sake of simplicity, we describe only a single motion detector unit, Fig 1.

In the simulations, we use a variety of motion detectors, which have a constant delay parameter, but have a range of different distances between the two photoreceptors in the single motion detector (d = x2-x1), Fig 1. The distances, d, vary from *d*_*min*_ *to d*_*max*_, (Table 1).

**Table 1:** Simulation’s parameters.

| Parameter | Value |
| --- | --- |
| A white pixel | 1 |
| A black pixel | 0 |
| A gray pixel | 0.5 |
| $d_{min}$ | 3 |
| $d_{max}$ | 15 |

### Directional selectivity (DS) motion detectors

Our model follows the anatomical and physiological findings in which the first layer of motion directionality responses is located in the T4 and T5 cells in the fly Lobula layer in the optical lobe; or in the starburst amacrine cells (SAC) in the mammalian retina (Alexander Borst and Helmstaedter 2015). The model applies the ability to detect motion in several directions as found in mammals and flies (Alexander Borst and Helmstaedter 2015).

We implemented the directional selectivity responses through Equations (5-8). The DS cells are divided into four types: 1) Rightward On DS cell 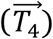, Equation (5). 2) Leftward On DS cell 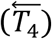, Equation (6). 3) Rightward Off DS cell 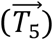, Equation (7). 2) Leftward Off DS cell 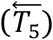, Equation (8). In order to calculate the direction of each pixel in Frame 1, Fig 2, we need to calculate the values of each DS type 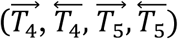 for each pixel. We calculated the responses by summing the element-wise multiplication (Hadamard product) of the pixels in Frame 1 with the shifted pixels in Frame 2, Fig 2, Equation (5-8). This type of calculation is equal to a directional correlation, i.e., we divided the full correlation to two separate correlation calculations: one for the positive direction (rightward, 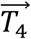) the other one for the negative direction (leftward, 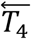).

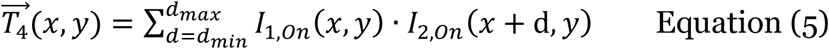

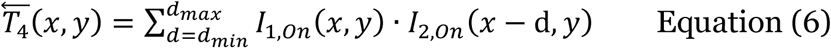

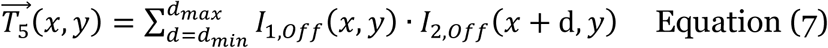

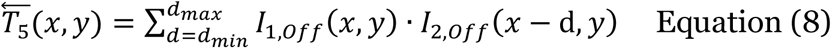

The equations (5-8) describe the correlation between the appearance of the first frame (Frame 1) and the presence of the second frame (Frame 2). The model also takes a secondary response (the disappearance of the first frame) into account, i.e., the rebound response results from the stimulus in the first frame. We, therefore, demonstrate the DS responses to this secondary response, as follows:

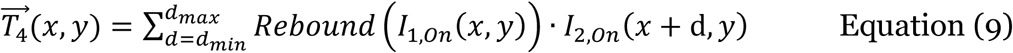

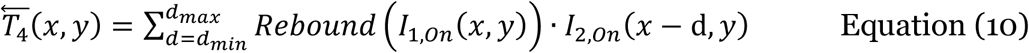

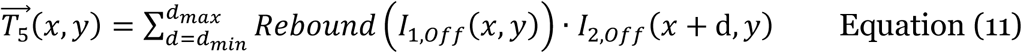

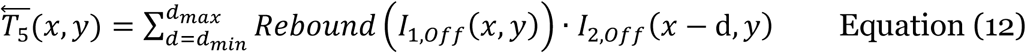

### Psychophysical Experiment

#### Stimuli

In the current experiment, we tested the two types of motions – phi and reverse-phi, with eight different apparent moving stimuli combinations, Table 1. These stimuli were adopted from Kitaoka (Kitaoka 2015). The moving stimuli are composed of a sequence of frames, where each frame in the sequence, Fig 4, is displayed for 100 ms. The objects in all frames are identical but rotated by about 6 pixels in the appropriate direction (clockwise or counterclockwise). This rotation is temporally adjusted to the temporal properties of the classical apparent motion.

**Fig 4:**
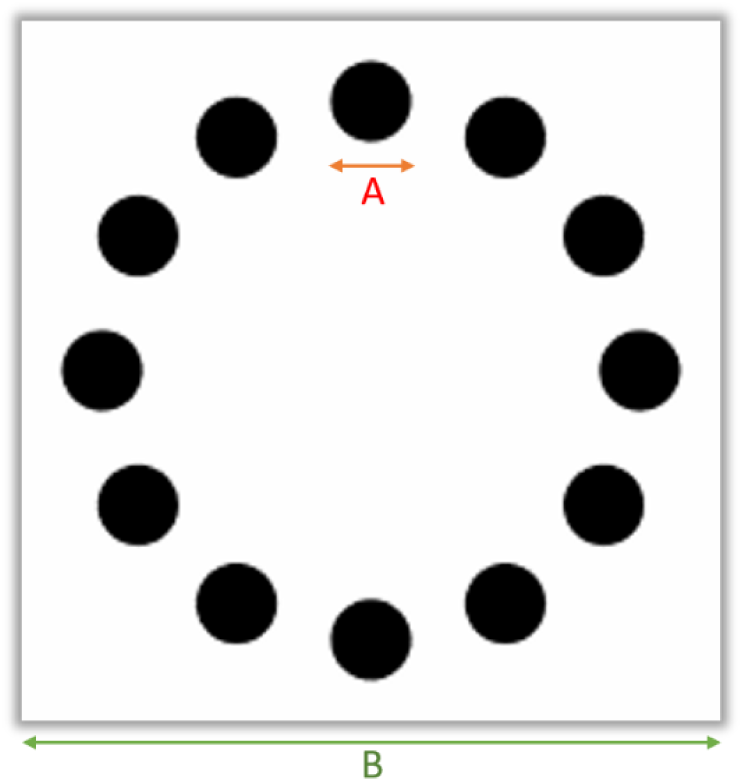
The basic stimulus of the psychophysical experiment consists of 12 circles. Each circle is 0.66^0^ visual degrees (A). The size of the stimulus is 5.75^0^ visual degrees (B). The luminance of the white background was 128 cd/mm, while the luminance of the black circles was 4.5 cd/mm, and the luminance of the gray background (beyond the size of the stimulus) was 34 cd/ mm (Table 2).

The eight stimuli, Table 1, are merged here into four types of stimuli: 1) Phi motion stimuli: two classical phi stimuli (with clockwise or counterclockwise rotation), where each is composed of 10 frames, Table 3: row 1-2,(Kitaoka 2015). 2) Reverse-phi motion stimuli: Two classical reverse-phi stimuli (clockwise or counterclockwise rotation), where each type of motion is composed of 10 frames. In these stimuli the even frames are displayed with an opposite (reverse) contrast, Table 3: row 3-4, (Kitaoka 2015). 3) Masked-phi motion stimuli: two types of masked phi stimuli (clockwise or counterclockwise rotation), which are built from the frames of the classical phi stimuli, but with the addition of a noise frame mask inserted between every two frames of the classical stimulus. The random mask is composed of uniformly distributed random noise, i.e., each pixel’s value is randomly selected from the range (0-1), for each noise frame, separately. This noise frame (mask) is displayed for a duration of 20 ms, Table 3: row 5-6. 4). Motion stimuli of masked reverse-phi: two types of masked reversed phi stimuli (clockwise or counterclockwise rotation), which are built from the frames of the classical reverse-phi stimuli. In this type of stimuli, we also added a mask frame, which displayed for 20 ms, between every two frames, similarly to the masked-phi stimuli Table 3: row 7-8. The total duration of the classical stimuli (the phi and the reverse-phi) presentation was 1000 ms. The total duration of the presentation of the masked stimuli was 1200 ms.

**Table 2:** The parameters of the experiment.

| Parameter | Value |
| --- | --- |
| Background Luminance (gray background) | 34 cd/ mm |
| white luminance | 128 cd/mm |
| Black luminance | 4.5 cd/mm |

**Table 3:** summarizes the stimuli conditions of the experiment. The experiment consists of 8 different videos; see supplementary videos. Each video presents 12 circles that have a real displacement (rotation) over the video’s different frames. The rotation direction is either clockwise or counterclockwise.

| No | Motion type | Mask | Real rotation |
| --- | --- | --- | --- |
| 1 | Phi | - | Clockwise |
| 2 | Phi | - | Counterclockwise |
| 3 | Reverse phi | - | Clockwise |
| 4 | Reverse phi | - | Counterclockwise |
| 5 | Phi | + | Clockwise |
| 6 | Phi | + | Counterclockwise |
| 7 | Reverse phi | + | Clockwise |
| 8 | Reverse phi | + | Counterclockwise |

#### Participants

Four adult observers, 3 males and one female, participated in the experiment. All participants had normal or corrected to normal vision.

#### Procedure

Observers viewed the stimuli on a flat-screen monitor placed at a distance of 62 cm. Participants registered a binary response to each trial, with participants pressing either of two keyboard buttons to indicate the perceived direction of motion (clockwise rotation or counterclockwise rotation). The observers, therefore performed force choice task. No feedback was provided.

#### Data Analysis

The observers’ results were assumed to represent a sample of a binomial distribution, due to the nature of the task. forced choice task. The proportion of correct responses (p = c/n), therefore, was computed from the ratio between the correct responses - (c) - (i.e. the perceived rotation direction is identical to the physical rotation direction) and the total number of group presentations (n), for each group. The standard deviations (SD) were calculated by using the binomial equation (referring to samples and not population) 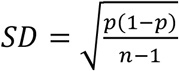 was calculated while considering the binomial distribution, and while this condition is satisfied when either p × n > 5 or (1 − p) × n> 5, depending on whether p is close to 1 or 0, (Wehrhahn 2006).

